# Phage toxin variants are linked to protection specificity in a defensive symbiont

**DOI:** 10.1101/2025.11.21.689672

**Authors:** Balig Panossian, Ailsa H. C. McLean, Vilas Patel, Taoping Wu, Muhammad Bilal Haider, Kerry M. Oliver, Lee M. Henry

**Affiliations:** School of Biological and Behavioural Sciences, Queen Mary University of London, London, United Kingdom; Department of Biology, University of Oxford, Oxford, United Kingdom; Department of Entomology, University of Georgia, Athens, Georgia, USA

**Keywords:** *Hamiltonella defensa*, symbiosis, host defence, parasitoid, genome evolution, bacteriophage

## Abstract

Insects often depend on symbiotic bacteria for protection, yet the mechanisms by which these microbes target specific natural enemies remain poorly understood. In aphids, different strains of the facultative symbiont *Hamiltonella defensa* provide highly specific protection against particular species of parasitoid wasps. To uncover the genetic basis of this specificity, we analyzed 26 *Hamiltonella* genomes and their toxin-encoding APSE bacteriophages with distinct protective phenotypes. Our analyses revealed that *Hamiltonella* strains share a conserved core genome but differ significantly in accessory gene content, reflecting their distinct evolutionary origins. Strikingly, we show that variation in toxin types is the key distinguishing feature of APSE phages in *Hamiltonella* strains that protect against different parasitoid species. These toxin repertoires include several novel candidates, such as variants with MAC/perforin domains and leucine-rich repeat (LRR) proteins previously unreported in insect defensive symbionts. We also reveal cases of multiple co-integrated APSE phages carrying different toxins within a single genomic locus. These findings implicate phage-borne toxins as primary determinants of enemy-specific defense and point to phage-driven toxin diversification as a major force shaping the functional evolution of this symbiosis. This work highlights how mobile genetic elements influence the ecological roles and diversification of protective symbionts.

## Introduction

Insects are attacked by a myriad of natural enemies, driving the evolution of diverse defensive strategies. In addition to developing resistance through the host genome, many insects rely on symbiotic bacteria to defend against attackers (Oliver et al. 2014). Defensive symbioses have now been identified across the Insecta and involve partnerships with diverse microbial taxa that offer protection against a broad range of natural enemies (Cornwallis et al. 2023). This includes microbes that protect against sterilization from nematodes, infection by viruses, and attack from fungal pathogens and parasitoid wasps (Scarborough et al. 2005; Hedges et al. 2008; Vorburger et al. 2010; Yadav et al. 2018). Given the widespread occurrence of these beneficial associations and their substantial contribution to host protection, defensive symbioses are now recognised as important factors influencing insect-natural enemy population ecology and coevolutionary dynamics (Vorburger and Perlman 2018; Monticelli et al. 2019). Moreover, symbionts can provide highly targeted resistance against specific natural enemies (Kellner and Dettner 1996; McLean and Godfray 2015; Mateos et al. 2016), yet our understanding of the genetic mechanisms underlying this variation is limited.

One well-studied defensive symbiont is the gammaproteobacterium *Hamiltonella defensa*, hereafter *Hamiltonella*, which is estimated to occur in around 40% of aphid species and also psyllids and whiteflies (Clark et al. 1992; Russell et al. 2003; Henry et al. 2015). *Hamiltonella* is predominantly vertically transmitted but can also be horizontally transferred between host lineages and even species (Sandström et al. 2001). Studies have shown that *Hamiltonella* is typically non-randomly distributed across aphid species and populations (Henry et al. 2022; Wu et al. 2022), and often occurs at high frequencies in host populations that carry the microbe. Although facultative, and therefore not essential for host survival, this symbiosis enhances host fitness by protecting against parasitoid wasps (Oliver et al. 2003). *Hamiltonella*’s defensive capabilities stem from its association with the bacteriophage APSE, which encodes eukaryotic toxins that target wasps (Van Der Wilk et al. 1999; Moran et al. 2005).

Experimental evidence has confirmed APSE’s role: phage loss eliminates protection (Oliver et al. 2009), while horizontal acquisition both in culture and *in vivo* restores the defensive phenotype (Brandt et al. 2017; Lynn-Bell et al. 2019). Homologous toxins in *Drosophila* also exhibit direct wasp-killing activity (Verster et al. 2023). Notably, *Hamiltonella* isolates vary in protection strength and specificity against different parasitoid wasp species (Oliver et al. 2005; Schmid et al. 2012; Cayetano and Vorburger 2013; Asplen et al. 2014; Martinez et al. 2014; Cayetano and Vorburger 2015; McLean and Godfray 2015; Martinez et al. 2016; Hopper et al. 2018; Wu et al. 2022) While specific toxin modules dictate the strength of protection against a single wasp species (Patel et al. 2023), the molecular basis of target specificity to different parasitoid species remains unclear.

The *Hamiltonella* strains found in European pea aphids (*Acyrthosiphon pisum*) provide striking examples of defensive variability (McLean and Godfray 2015; Leclair et al. 2016). In Europe, pea aphids form a complex of genetically differentiated, plant-associated biotypes (Peccoud et al. 2009a), with populations on *Medicago sativa*, *Lotus pedunculatus*, and *Ononis spp*. (hereafter *Medicago*, *Lotus*, and *Ononis*) each predominantly harboring distinct *Hamiltonella* strains at high frequencies (Henry et al. 2013). McLean and Godfray (2015) found intriguing variation in the defensive properties of these strains – *Hamiltonella* from *Medicago-*associated pea aphids provided strong protection against the braconid parasitoid, *Aphidius ervi*, but not the chalcid parasitoid *Aphelinus abdominalis*, whereas *Hamiltonella* from *Lotus*-associated aphids protected against *A. abdominalis*, but not *A. ervi*. In contrast, the *Hamiltonella* from *Ononis-A. pisum* offered no protection against either of these natural enemies. A study by Wu et al (2022) found a similar pattern in that the *Hamiltonella* strains from *Macrosiphoniella artemisiae* and *Medicago* pea aphids each protected against its main parasitoid species (*Aphidius absinthii* and *A. ervi,* respectively), but not the other. Despite these striking differences in defensive capabilities, the genetic basis of this target specificity remains unknown. Here, we hypothesize that toxins encoded by APSE phage variants mediate parasitoid-specific defense.

In this study, we examined 26 *Hamiltonella* genomes, including 13 with known protective phenotypes, to explore the genomic basis of their defensive variation. This included 10 newly sequenced strains from the European pea aphid biotypes associated with *Medicago*, *Lotus*, and *Ononis* biotypes, two strains from *M. artemisiae* and *Macrosiphum euphorbiae* (Wu et al. 2022), and 16 publicly available genomes. Our genome analysis reveals that toxin modules are the key structural difference among APSE strains, with several novel toxins identified; our results suggest these variants underlie the strain-specific protection *Hamiltonella defensa* confers against different parasitoid wasps.

## Results

### Evolutionary origins of the Hamiltonella strains investigated in our study

We determined the phylogenetic placement of *Hamiltonella* strains included in our study by constructing a maximum likelihood (ML) phylogenetic tree based on 147 single-copy orthologous genes that included all published *Hamiltonella* genomes where it is a facultative symbiont. This produced a well-supported tree that notably places the *Hamiltonella* from the *Ononis* biotype of *A. pisum* and *Hamiltonella* from *M. artemisiae* as the most basal clades in the *Hamiltonella* phylogeny (Fig. 1). Although this placement is likely influenced by sampling limitations - most samples are from *A. pisum* - it demonstrates the *Hamiltonella* clade associated with *Ononis*-*A. pisum* and *M. artemisiae* are ancestral to other known lineages found in aphids. As shown in previous studies, we confirm that lineages from the *Medicago* biotype are paraphyletic (Henry et al. 2013; Guyomar et al. 2018) (Fig. 1). *Hamiltonella* from *Lotus* biotype *A. pisum* and *M. euphorbiae* aphids form a clade that includes several strains associated with the *Medicago* biotype (Fig. 1). Our results also confirm that *Medicago*-associated strains tend to form two major clades, with representatives from both found in the US and the UK. Interestingly, *Hamiltonella* lineages that belong to the largest clade associated with *A. pisum* on *Medicago* are closely related to several lineages collected from *Cinara* conifer aphids in France and Korea, as well as the social aphid *Ceratovacuna* in Japan. This demonstrates that *Hamiltonella* has been horizontally transferred across distantly related host species (Russell et al. 2003; Guyomar et al. 2018), and suggests that the pea aphid biotypes likely acquired their *Hamiltonella* strains from other aphid species after their post-Pleistocene radiation to different host plants (Peccoud et al. 2009b).

**Figure 1.**
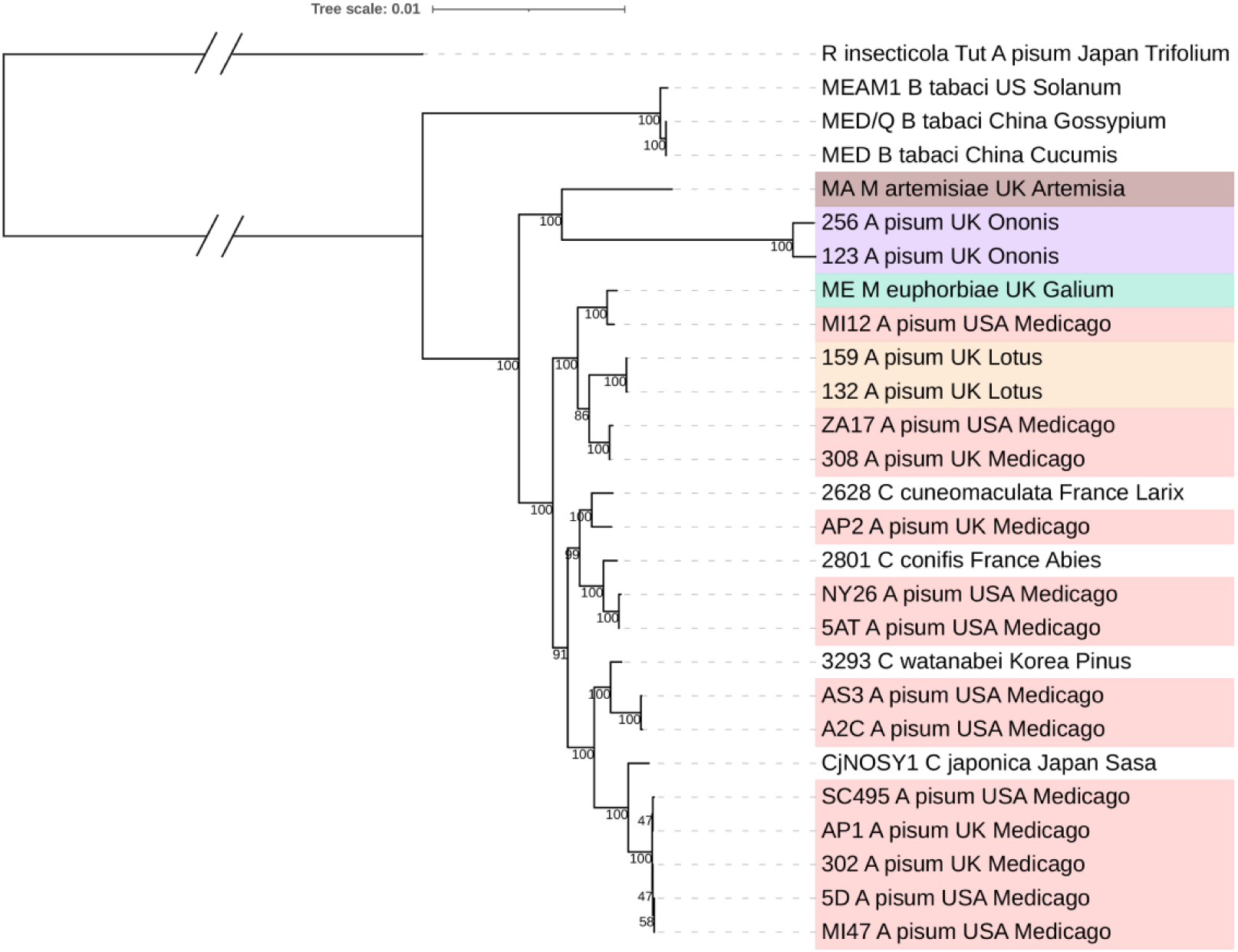
Maximum-likelihood phylogenetic tree of facultative *Hamiltonella* strains. based on 147 housekeeping genes rooted to *R. insecticola*. Values at nodes reflect bootstrap support. Colours represent the plant association of each *Hamiltonella* strain; *M. artemisiae* in Brown, *M. Euphorbia* in Green, and *A. pisum* biotypes *Ononis* in Purple, *Medicago* in Pink, and *Lotus* in Orange.

### Genome characteristics of aphid biotype- and species-associated Hamiltonella strains

We conducted detailed analyses on 19 *Hamiltonella* genomes from two sources. First, we sequenced 10 strains commonly associated with five aphid lineages in Europe—including *Lotus*, *Ononis*, and *Medicago* biotypes of the pea aphid (*A. pisum*), as well as the potato aphid (*M. euphorbiae*) and mugwort aphid (*M. artemisiae*), many of which have known protective phenotypes (Table 1). We then included 9 previously sequenced *Hamiltonella* genomes from pea aphids collected on *Medicago* in the USA (Table 1, Table S1), some of which also have known defensive properties. We included *Hamiltonella* strains from *Cinara* and *Ceratovacuna* aphids and *Bemisia* whiteflies only in the phylogeny, given the limited experimental evidence for parasitoid-wasp protection. We found that *Hamiltonella* from *A. pisum* on *Ononis* (256, 123) have significantly smaller genomes, 1.845±0.035Mb, than strains from the *Medicago* biotype at 2.17±0.032Mb (Padj=0.006**), and the *Hamiltonella* from *M. artemisiae* (Ma) is also reduced in size at 1.94Mb. The reduced size of the *Ononis* and MA *Hamiltonella* strains suggests that these more ancestral lineages have undergone significant genome rearrangements and may have experienced erosion over their evolutionary history (McCutcheon and Moran 2012). *Medicago*-associated *Hamiltonella* strains from Europe had largely similar genome characteristics to those from the USA (Table 1, Supplementary Table S2). Phage and plasmid numbers were found to be variable across strains and biotypes. Although most genomes had >99% completeness and >100x coverage, we could not confirm the presence or absence of plasmids in all *Hamiltonella* strains (N/A’s) due to sequencing with short read Illumina technology in some cases (Table S2).

**Table 1.** Metadata describing the *Hamiltonella* strains analyzed in this study. Protective phenotype describes the species of parasitoid wasp the *Hamiltonella* strain was tested against and whether it provided protection or not.

| Organism Qualifier | Geographic Location | Plant | Protective Phenotype | Genome size (Mb) | Plasmids | Intact Phages | Genome |
| --- | --- | --- | --- | --- | --- | --- | --- |
| 5AT | USA | Medicago | Protection *A. ervi* (Moran et al. 2005) | 2.169 | 1 | 2 | Chevignon et al. 2018 |
| A2C | USA | Medicago | No Protection (phage lost) (Oliver et al. 2009) | 2.204 | 3 | 2 | Chevignon et al. 2018 |
| AS3 | USA | Medicago | Protection *A. ervi* (Oliver et al. 2009) | 2.262 | 2 | 3 | Chevignon et al. 2018 |
| ZA17 | USA | Medicago | N/A | 2.273 | 3 | 1 | Chevignon et al. 2018 |
| NY26 | USA | Medicago | N/A | 2.122 | 0 | 3 | Chevignon et al. 2018 |
| MI47 | USA | Medicago | N/A | 2.276 | 5 | 3 | Boyd et al. 2021 |
| 5D | USA | Medicago | N/A | 2.266 | 5 | 2 | Boyd et al. 2021 |
| MI12 | USA | Medicago | N/A | 2.253 | 2 | 1 | Boyd et al. 2021 |
| SC_495 | USA | Medicago | N/A | 1.949 | N/A | 1 | Blow et al. 2020 |
| 132 | UK | Lotus | Protection *A. abdominalis*<br>No protection *A. ervi* (McLean et al. 2015) | 2.08 | 4 | 4 | This study |
| 159 | UK | Lotus | Protection *A. abdominalis*<br>No protection *A. ervi* (McLean et al. 2015) | 2.05 | 3 | 3 | This study |
| 256 | UK | Ononis | No Protection (McLean et al. 2015) | 1.81 | N/A | 2 | This study |
| 123 | UK | Ononis | No Protection (McLean et al. 2015) | 1.88 | N/A | 3 | This study |
| 308 | UK | Medicago | Protection *A. ervi*<br>No protection *A. abdominalis* (McLean et al. 2015) | 2.29 | 4 | 6 | This study |
| 302 | UK | Medicago | Protection *A. ervi*<br>No protection *A. abdominalis* (McLean et al. 2015) | 2.37 | 5 | 3 | This study |
| MA | UK | Artemisia | Protection *A. absinthii*<br>No protection *A. ervi* (Wu et al. 2022) | 1.94 | N/A | 2 | This study |
| ME | UK | Galium | No protection *A. absinthii*, *A. rhopalosiphii*, *A. ervi* (Wu et al. 2022) | 2.16 | N/A | 2 | This study |
| AP1 | UK | Medicago | Protection *A. rhopalosiphii*, *A. ervi* (Wu et al. 2022) | 2.12 | N/A | 3 | This study |
| AP2 | UK | Medicago | No protection *A. rhopalosiphii*<br>Protection *A. ervi* (Wu et al. 2022) | 2.03 | N/A | 4 | This study |

### Gene content of European Hamiltonella strains differ in several functional categories

Our pangenome analysis of *Hamiltonella* revealed that strains share a large core genome but have notable differences in their accessory genomes (Fig. 2A&B). Only 12 and 16 genes were unique to *Hamiltonella* from each of the *A. pisum Lotus* biotype and *M. euphorbia* from *Galium* sp. respectively, and no genes were uniquely present in the *Hamiltonella* from the *Medicago* biotype. In contrast, the *Ononis* biotype and *M. artemisiae* from *Artemisia* sp. associated *Hamiltonella* had the greatest gene content divergence containing respectively 109 and 215 genes only found in these strains (Fig. 2A).

**Figure 2.**
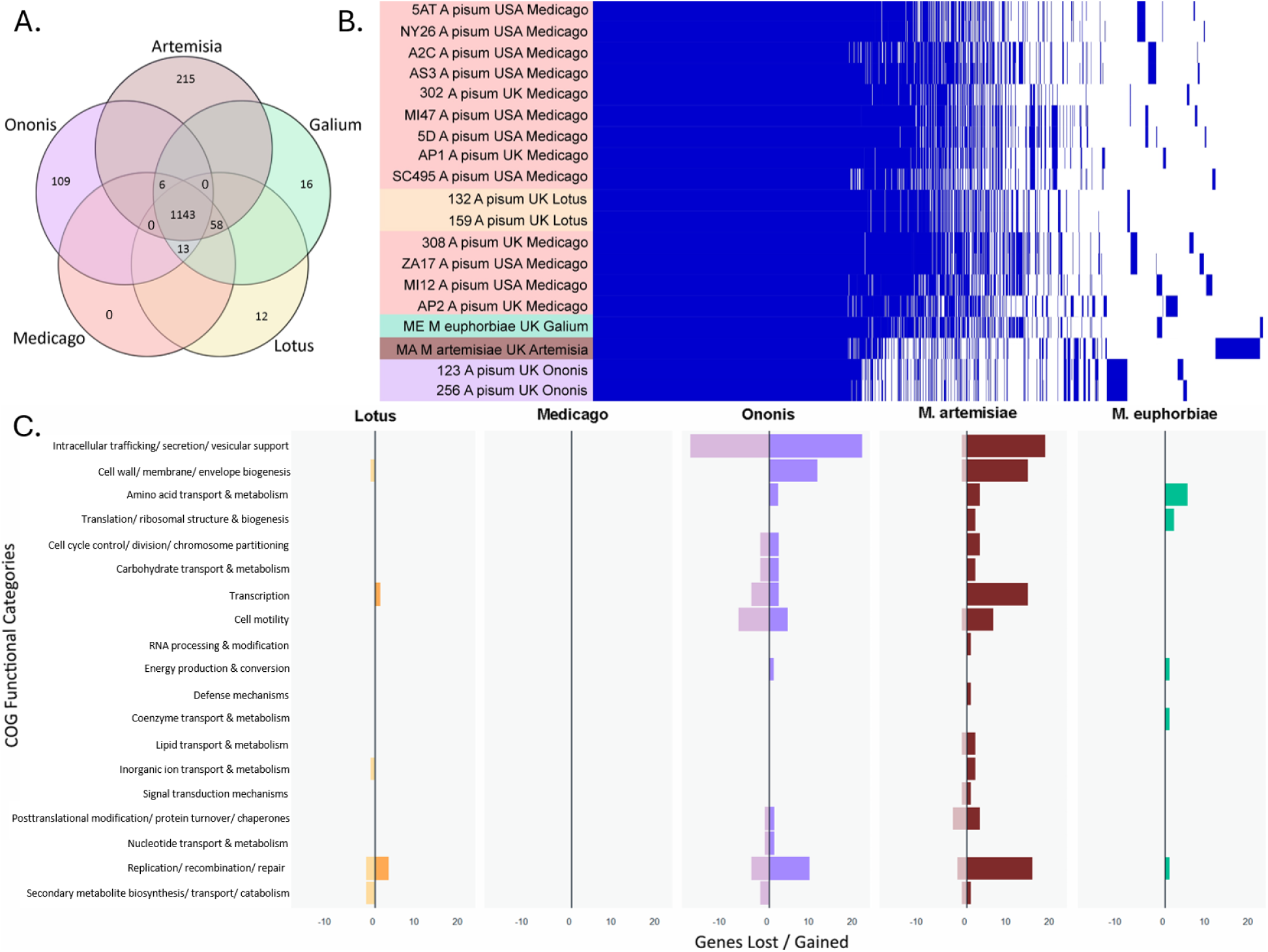
*Hamiltonella* genome comparisons. **(A)** Gene content distribution among the *Hamiltonella* genomes analyzed showing the number of shared/unique genes across the three European *A. pisum* biotypes (*Ononis*, *Medicago* and *Lotus*) and *M. euphorbiae* collected from *Galium aparine* and *M. artemisiae* from *Artemisia vulgaris*.; **(B)** Pangenome matrix of the *Hamiltonella* genomes illustrates the conserved core genome and highlights the unique accessory genome for samples of the five different biotypes. Colours represent *Hamiltonella* from *M. artemisiae* (brown), *M. euphorbiae* (green), and the *A. pisum Ononis* (purple), *Medicago* (pink) and *Lotus* (orange) biotypes. **(C)** Functional categories of genes uniquely present/absent in the genomes of the ten new *Hamiltonella* genomes (two from each pea aphid biotype, one each from *M. euphorbiae* and *M. artemisiae*) from this study.

All the genes uniquely present in the *Hamiltonella* from the *Lotus* biotype were APSE phage genes (Supplementary Table S3). This strongly implicates APSE variation as the driver of the contrasting defensive phenotypes.

A functional comparative analysis of European *Hamiltonella* genomes, based on Clusters of Orthologous Groups (COGs), revealed the greatest variability in three functional categories: (i) Intracellular trafficking, secretion, and vesicular support (including secretion system machinery), (ii) Replication, recombination, and repair, and (iii) Cell wall/membrane/envelope biogenesis. These differences were particularly pronounced in *Hamiltonella* from the *A. pisum Ononis* biotype and from *M. artemisiae* (Fig. 2C).

Large differences in the number of genes involved in Intracellular trafficking, secretion system machinery, and vesicular support (COG category U; Table S3) were attributed to the *A. pisum Ononis Hamiltonella* losing genes that encode the Type II/IV secretion systems. Notably, *Ononis* strains were devoid of all 13 genes required to construct and assemble a functional Type 2 Secretion System (T2SS) as well as 18 genes required for the Type 4aP secretion system (T4aPSS) (Fig. S1). Loss of secretion systems, especially T2SS, may affect pathogenicity and within-host fitness in *Ononis Hamiltonella*, as these are key genetic factors underlying these traits in free-living *Yersinia* relatives (von Tils et al. 2012). In contrast, the genes gained by *Hamiltonella* from *M. artemisia* almost entirely coded for intracellular transporters (Table S3).

*Hamiltonella* strains also carried unique sets of genes involved in Replication, recombination and repair (COG category L; Table S3). These include bacteriophage mobilization genes, integrases, resolvases, and transposases. Genes uniquely missing from the *Ononis Hamiltonella* strains include a DNA repair gene *phrB* coding for a photolyase. Genes influencing Cell wall, membrane and envelope biogenesis (COG category M; Table S3) uniquely gained in the *Ononis Hamiltonella* include *eaeH*, which mediates attachment to eukaryotic cells (Sheikh et al. 2014), as well as genes controlling cell (*rfbF, rfbG, fcl*) (Zhang et al. 1993) and colony morphology (*ddhC*) (Anriany et al. 2006). *Ononis Hamiltonella* were uniquely missing the *licD* gene that alters the cell surface glycans to promote adherence and invasion. In contrast, *Hamiltonella* from *M. artemisia* uniquely carries the *licA*, *licC* and *licD* genes, which are involved in eukaryotic mimicry and colonization (Zhang et al. 1999), and the *iabB* gene, which encodes for an invasion protein (Klein et al. 2000), and the adhesion gene *pmp10*. The *prgK* surface lipoprotein coding gene was also uniquely absent from the MA strain. *Hamiltonella* strains from the *A.pisum Medicago* and *Lotus* biotypes, and *M. euphorbiae*, did not show any unique gains and losses in genes coding for cell membrane structure. Together, these results suggest *Hamiltonella* strains differ in genes functionally important for interacting with hosts, which may contribute to host-symbiont incompatibility (Łukasik et al. 2015; Wu et al. 2022). Additionally, the *Hamiltonella* strains from *M. artemisiae* and *Ononis* pea aphids had a relatively higher number of cellular motility genes that were similar to bacterial flagella. Transcription factors were also detected at higher copy numbers in the *M. artemisiae* strain, some of which were multiple copies of core genes, which could be a result of duplication events. A full list of genes gained and lost, including a detailed annotation, can be found in the supplementary material (Table S3).

Our genome analysis also revealed that the APSE phage was absent from the *Hamiltonella* ME strain from *M. euphorbiae* (Wu et al. 2022). As *Hamiltonella* can lose APSE upon transfer to laboratory colonies (Oliver et al. 2009; Lynn-Bell et al. 2019), this likely explains the ME strain’s lack of protection reported previously (Wu et al. 2022).

### Variation in APSE phage toxin genes correlates with protective phenotypes

We investigated the genetic basis of parasitoid-species–specific resistance in European *Hamiltonella* by comparing the ∼40-kb gene content of their associated APSE bacteriophages, key mediators of parasitoid defence. We found that while the replication and virion assembly modules of APSE phages are highly conserved, their toxin-encoding "module 3" exhibits striking diversity (Fig. 3), consistent with previous reports (Rouïl et al. 2020; Boyd et al. 2021).

**Figure 3.**
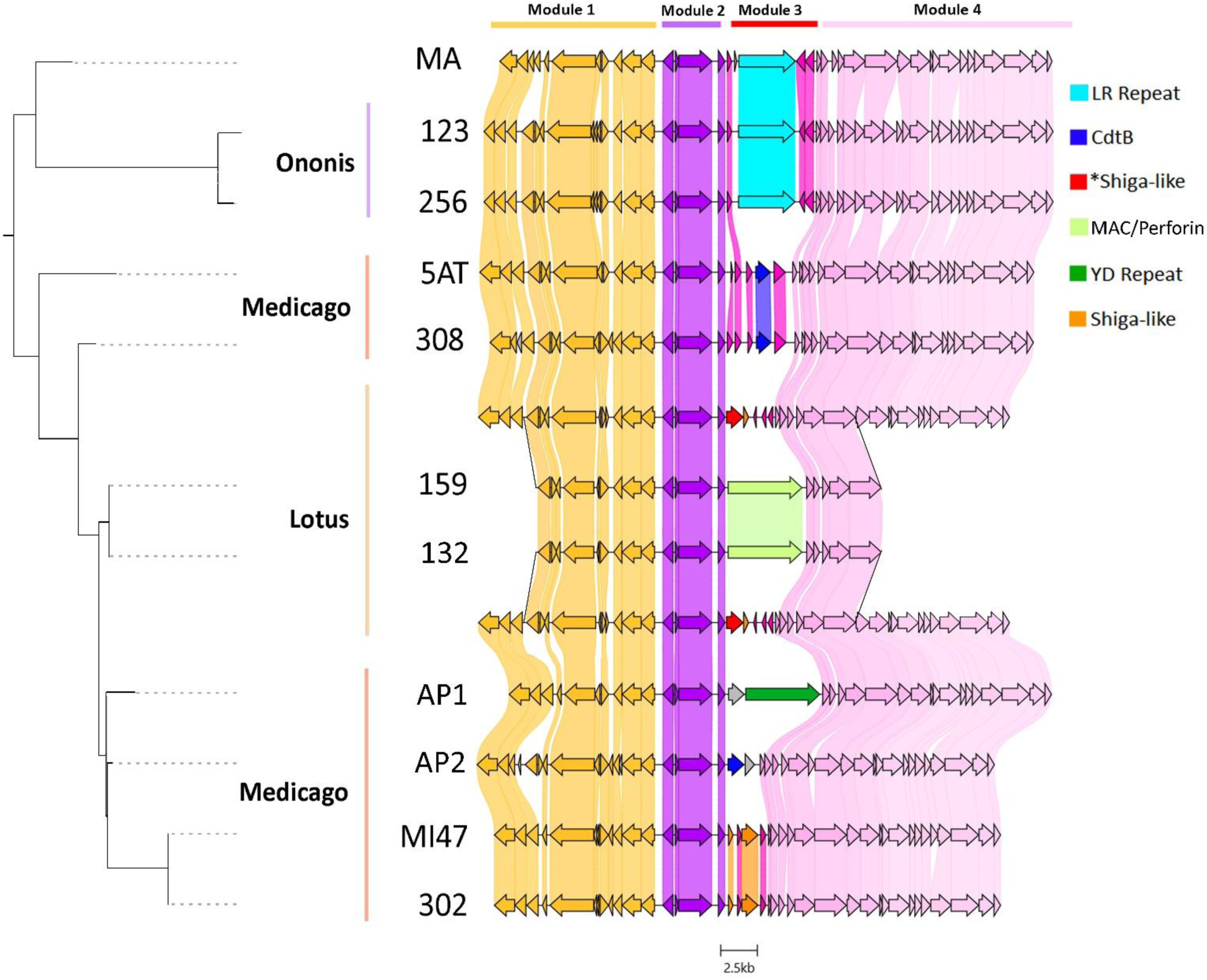
- **Comparison of APSE bacteriophage genomes** from the newly sequenced European *Hamiltonella* genomes. Previously sequenced APSE strains 5AT and MI47 are included for comparison. Host plant (biotype) origins of each strain is indicated on the left and APSE modules are noted above. Note that most structural variation of the genomes occurs in Module 3, which contains genes encoding for eukaryotic toxins. In samples 132 and 159 from *Lotus* pea aphids, the genes of the two APSE phages are illustrated as two branches stemming from a shared APSE backbone, indicating the multiple copies of backbone genes and unique toxin-bearing modules integrated at the same genomic locus (Fig. S2).

Most notably, two *Hamiltonella* strains from the *Lotus* biotype (159 and 132) - which confer protection against the chalcid parasitoid *Aphelinus abdominalis* - harbour a unique APSE genotype composed of multiple, co-integrated APSE phages within a single genomic locus (Fig. 3, Fig. S2). Each of these co-resident phages carries a distinct, previously uncharacterized toxin module. One encodes a novel variant of the previously characterised Shiga-like toxin (377 Amino Acids, AAs) in APSE-1 (Fig. S3A), but with the alpha and beta subunit genes arranged in reverse order and followed by three small transposases (Fig. 3). The second harbours a large (1678 AA) gene containing a membrane attack complex/perforin (MAC/Perforin) domain, marking the first report of such a toxin in *Hamiltonella* (Fig. 3, Fig. S3). Analysis of sequence data suggests these two APSEs are integrated within the exact same genetic locus in both strains (132, 159) of the *Hamiltonella* genomes from *Lotus* biotype pea aphids. Although the precise orientation of the two toxin loci remains unresolved, haplotype-phased assembly graphs suggest the two APSE phages share partial peripheral backbone Modules 1 and 4 and an identical, fully duplicated Module 2, after which each carries a distinct Module 3 encoding different toxin genes (Fig. 3, Fig. S2).

In *Hamiltonella* from the *Medicago* biotype, which protects against the braconid *A. ervi*, we found two toxin types: one phage encodes a cytolethal distending toxin B (*cdtB*; strain 308), while another (strain 302) carries a divergent Shiga-like toxin distinct from that in the *Lotus* biotype. In contrast, the *Hamiltonella* strain from *M. artemisiae* (MA), which protects specifically against *A. absinthii* but not related *A. ervi* (Wu et al. 2022), contains an APSE phage with a large (1285 AA), highly divergent gene encoding a Leucine-Rich Repeat (LRR) domain protein (Fig. S3B). Similar LRR toxins were also found in APSE phages from the *Ononis* biotype (strains 123 and 256), with 97% nucleotide and 94% amino acid identity to the MA variant, suggesting phylogenetically related phage strains tend to carry similar toxin families (Fig. 3).

Phylogeny on the left is a pruned version of the conserved core APSE backbone from supplementary Fig. S4.

## Discussion

Our results show that variation in mobile genetic elements, specifically APSE bacteriophages and their toxin modules, likely determines resistance specificity against different natural enemies. This suggests a dynamic interplay between *Hamiltonella* and different APSE phage isolates underlies variation in host protective phenotypes.

### APSE Phage toxin variation is associated with target-specific symbiont defense

Previous studies show that *Hamiltonella* often protects aphids specifically against the parasitoids that most frequently attack them in nature (Łukasik et al. 2015; McLean and Godfray 2015; Martinez et al. 2016; Wu et al. 2022). Our genomic analysis confirms that APSE phage backbones are highly conserved, whereas their toxin modules are hotspots of variation that correlate with parasitoid-specific protection (Boyd et al. 2021). We identified several novel toxin candidates, most notably a large protein containing a membrane attack complex/perforin (MAC/perforin) domain in the APSE phages of Lotus-associated *Hamiltonella*. This is the first report of such a toxin in *Hamiltonella* or its APSE phage. In other systems, including eukaryotic immunity and bacterial pathogenesis, MAC/perforin domain proteins function by oligomerizing to form pores in target cell membranes, leading to cell lysis (Dunstone and Tweten 2012). The presence of this previously unknown toxin class in APSE phages hints at a novel mechanism for parasitoid wasp resistance.

These same APSEs, which protect against the chalcid *A. abdominalis*, also encode a divergent Shiga-like toxin; while other APSEs carry Shiga-like toxins that target braconids such as *A. ervi* (Asplen et al. 2014; McLean and Godfray 2015). A highly conserved Shiga-toxin β subunit suggests similar binding to Gb3-like receptors, whereas divergence in the α subunit could alter ribosomal injury and thus host specificity (Melton-Celsa 2014). Resistance to *A. abdominalis* may therefore arise from the MAC/perforin toxin, the distinct Shiga-like variant, or their synergy. Regardless, these findings suggest that both the acquisition of new toxin families and the sequence divergence of existing ones are potential evolutionary routes driving diversification of protective phenotypes in defensive symbioses.

Our genome comparison also revealed a LRR putative toxin gene in the APSEs of the *Hamiltonella* from *M. artemisiae* (MA) and *Ononis* pea-aphid biotypes. Despite having no known prokaryotic orthologue, its placement within the APSE toxin module, combined with the known virulence of LRR proteins in *Yersinia* (Hines et al. 2001), implicates it in parasitoid defense. Consistent with high specificity, MA *Hamiltonella* protects *M. artemisiae* against the parasitoid *A. absinthii* but not *A. ervi* (Wu et al. 2022). Interestingly, the APSE phage in *Ononis Hamiltonella*, which is nearly identical to the one found in *M. artemisiae*, also does not protect against *A. ervi,* or *A. abdominalis* (McLean and Godfray 2015). This similarity suggests the toxin in the *Ononis* APSE may target an untested parasitoid. Ononis pea aphids are attacked by *A. aedyi* (Starý 2006), and parasitoid communities strongly predict *Hamiltonella* genotype distributions (Wu et al. 2025), and potentially the APSE and toxins they carry. Future research should test whether the LRR gene found in *Ononis* APSE provides specific protection against *A. aedyi* and assess its activity against other parasitoids.

### APSE Phage co-integration as a novel form of protection diversification

In *Lotus A. pisum* (clones 132 and 159), the APSE toxin module is duplicated, producing two module-3 copies that encode different toxins (Fig. S2). Our assemblies indicate these APSEs co-integrate at the same *Hamiltonella* locus - a rare genotype potentially driven by recombination at module boundaries (Rouïl et al. 2020). Although this is the first evidence of co-integration occurring in *Hamiltonella*, analogous co-integration of Shiga-toxin carrying prophage does occur in *E. coli* (Nakamura et al. 2021). Such parallel co-integration could rapidly expand a symbiont’s defensive arsenal by combining toxins rather than adding them sequentially. This highlights how modular, mobile phages can help generate multi-component protective phenotypes in the host-parasitoid arms race. Thus, we have shown that APSEs in European *Hamiltonella* diversify not only via toxin-module variation but also through parallel co-integration within a single genome - a previously unrecognized route to enhanced defense.

## Conclusion

Mobile genetic elements (MGEs), particularly bacteriophages, are major engines of functional innovation in symbiotic bacteria (Hosokawa et al. 2016; Gerth et al. 2021; Siozios et al. 2024). Our work underscores the central role of MGEs, specifically bacteriophages, as drivers of functional innovation in defensive symbionts. We demonstrate that phage-encoded toxin repertoires are the primary determinants of target-specific protection in the *Hamiltonella*-aphid symbiosis, which evolve by both sequence divergence and the acquisition of novel families such as the MAC/perforin protein. We also show that phages can co-integrate within a single host genome, enabling the rapid assembly of multi-component defences. Together, these findings reveal how the modularity and mobility of phage genomes generate the variation and specificity needed for symbiont-mediated resistance, driving adaptation in the coevolutionary arms race between insects and their natural enemies.

## Materials and Methods

### Sample collection and genome sequencing

Ten aphid clones carrying *Hamiltonella* with distinct defensive phenotypes were included in this study (Table 1). Six pea aphid clones from *Lotus pedunculatus, Ononis spinosa, and Medicago sativa* were obtained from laboratory cultures at the University of Oxford (McLean and Godfray 2015). Potato aphid (*M. euphorbiae*), mugwort aphid (*M. artemisiae*), and two pea aphid lineages were clones maintained from a previous study in our lab (Wu et al. 2022). We attempted to *in vitro* culture all pea aphid *Hamiltonella* strains established through the methodology developed by Brandt et al (2017), however, only strains from *Lotus* and *Medicago* were successfully grown using this method. Sufficient concentrations of pure bacterial pellets were recovered, and DNA was extracted as described in (Weldon et al. 2013). Purified *Hamiltonella* genomes were sequenced on a PacBio Sequel II platform at an average yield of 88,851 x 9300bp reads per sample. Additional strains, including those that were unculturable, were sequenced using short-read Illumina technology. DNA was extracted from individual *Ononis, M. euphorbiae,* and *M. artemisiae* aphids using the EZNA Insect DNA kit (Omega Bio-Tek, Norcross, Ga, USA) and they were sequenced on an Illumina NovaSeq S4 following the NEBNext Ultra II FS kit library prep protocol to generate paired-end reads of 2x150bp length at an average of 200 million reads per sample. All samples were sequenced at the Centre for Genomic Research at the University of Liverpool.

### Genome assembly and quality control

Sequencing reads were checked for quality and adapter content using FastQC (Simon 2010) and MultiQC (Ewels et al. 2016), and were trimmed using FastP (Chen et al. 2018) with default parameters. Long reads were assembled using MetaFLYE v2.9 (Kolmogorov et al. 2020) with parameters --pacbio_hifi and 5 polishing iterations. Short reads were assembled in two steps. First, SPAdes v3.15.4 (Bankevich et al. 2012) was used in assembly only mode with default parameters. The reads were mapped to the assembly using BWA-mem (Li and Durbin 2009) and taxonomic identities were assigned using megablast (Morgulis et al. 2008) and DIAMOND (Buchfink et al. 2015) searches against the NCBI’s non-redundant Refseq nucleotide and protein databases respectively. These were examined using BlobTools v1.1.1 (Laetsch and Blaxter 2017) where contigs matching *Hamiltonella* showing a distinct coverage and GC content were extracted (Fig. S5). Then, *Hamiltonella* reads were extracted using SAMtools (Li et al. 2009) and reassembled to give more contiguous and complete genomes with SPAdes v3.15.4 (Bankevich et al. 2012) in careful mode using error correction and kmer sizes of 33, 55, 77, 99, and 127.

All final assembly graphs were manually inspected in Bandage for uniform coverage and contiguity, and Blastn (Camacho et al. 2009) searches against the NCBI nonredundant nucleotide database were done to validate the taxonomic identities. Assembly completeness and contamination was assessed with CheckM (Parks et al. 2015) using f_*Enterobacteriaceae* as the marker lineage that has 546 genomic marker genes. Genomic markers and functional genes that were automatically assigned as missing were verified using HMM and blastp searches of the target genes on the unfiltered metagenome contigs.

### Phylogenetics

To determine the phylogenetic placement of the *Hamiltonella*s investigated in our study, we generated a phylogeny of all facultative *Hamiltonella*s with published genomes downloaded from Genbank. The GToTree v1.8.2 (Lee 2019) pipeline was run in default settings with 1000 bootstraps in RAxML v8 (Stamatakis 2014) on a concatenated alignment of the 172 orthologous genes universally present in Gammaproteobacteria. The phylogeny was visualized and rooted to *Regiella insecticola* Tut (GCA_013373955.1) using ITOL (Letunic and Bork 2021).

Phylogenies of APSE phages were generated by identifying the core genes shared by all APSE phage backbones using Orthofinder v2.5.4 (Emms and Kelly 2019). Orthologues were aligned and concatenated using Muscle 5.1.linux64 (Edgar 2022) and TrimAL v1.4.rev15 (Capella-Gutiérrez et al. 2009). PhyML v3.0 (Guindon et al. 2010) was run with default settings and automatic model selection and 500 bootstraps to generate the phylogeny which was visualized and rooted to the APSEs from *Hamiltonella* in *Bemisia tabaci*.

To determine the phylogenetic placement of the newly identified shiga-like toxin alpha subunit, the Lucine-rich repeat and MAC/Perforin domain containing toxin genes, we identified 100 of each of their closest hits on Genbank which were aligned using Muscle 5.1.linux64 (Edgar 2022). The alignment was used in PhyML (Guindon et al. 2010) to generate a phylogenetic tree with default parameters and automatic model selection with 500 bootstraps. The tree was midpoint rooted, pruned, and visualized in ITOL (Letunic and Bork 2021).

### Annotation and pan-genome analysis

*Hamiltonella* genomes were annotated using Prokka v1.14.6 (Seemann 2014) with default settings in --compliant mode. Genomes downloaded from Genbank were re-annotated to ensure uniformity. Roary v3.13 (Page et al. 2015) was used with -e -i 90 -s settings to identify core genes common at (90% amino acid identity cutoff) in all genomes and accessory genes unique to *Hamiltonella* strains of each biotype. The pangenome matrix was visualized using Phandango (Hadfield et al. 2018). Genes uniquely present and absent in *Hamiltonella* from each biotype were annotated with COG, KEGG, GO terms, and Pfam domains using eggnog v6.0 (Hernández-Plaza et al. 2023). Visualization of the COG terms was done with GGplot2 in R v4.3.2 (Ihaka and Gentleman 1996).

### Comparative analysis of gene function

Pan-genome analysis revealed major gene rearrangements belonged to three main functional groups – i) Intracellular trafficking, secretion, and vesicular support, ii) Replication, recombination and repair, and iii) Cell wall/membrane/envelope biogenesis genes. After annotating the *Hamiltonella* genomes we identified major functional groups of genes gained and lost in different lineages, with a particular focus on the functional categories where major gains/losses have occurred. We used Phaster (Arndt et al. 2016) to identify and extract bacteriophage sequences from the assemblies. It was run using default parameters, and bacteriophage sequences that were flagged as incomplete were filtered out. KEGG mapper (Kanehisa and Sato 2020) was used to check the presence of genes involved in metabolic pathways for amino acids, vitamins, carbon, lipids, and nucleotide metabolism and transporter genes. The TXSscan HMM model was used through MacSyFinder v2 (Abby and Rocha 2017) using the unordered replicon parameter to search for genes coding for bacterial secretion system machinery.

### APSE phage genome reconstruction

Bacteriophage contigs with annotated hits to APSE phages from the Phaster (Arndt et al. 2016) results were identified and flagged, and genome sequencing reads matching those contigs were reassembled with strict parameters as described above to yield contiguous APSE sequences. The initial assemblies were also checked for APSE sequences using Blastn searches of core APSE genes. The APSE genes were aligned and visualized using Clinker (Gilchrist and Chooi 2021).

### Statistics

Genome sizes were grouped by parent aphid species (and biotype for pea aphids) carrying each *Hamiltonella* strain. Standard errors were calculated for each group, and we used Levene’s to test for variance homogeneity. ANOVA tested genome size differences among categories, followed by Tukey’s HSD for post-hoc comparisons. Analyses were conducted in R v4.3.2 (Ihaka and Gentleman 1996) using dplyr, tidyr, and car packages.

## Data availability

The genome assemblies generated in this study have been deposited in the NCBI GenBank database under BioProject accession number PRJNA1338369.

