## Supplementary figures for "Phage toxin variants are linked to protection specificity in a defensive symbiont"

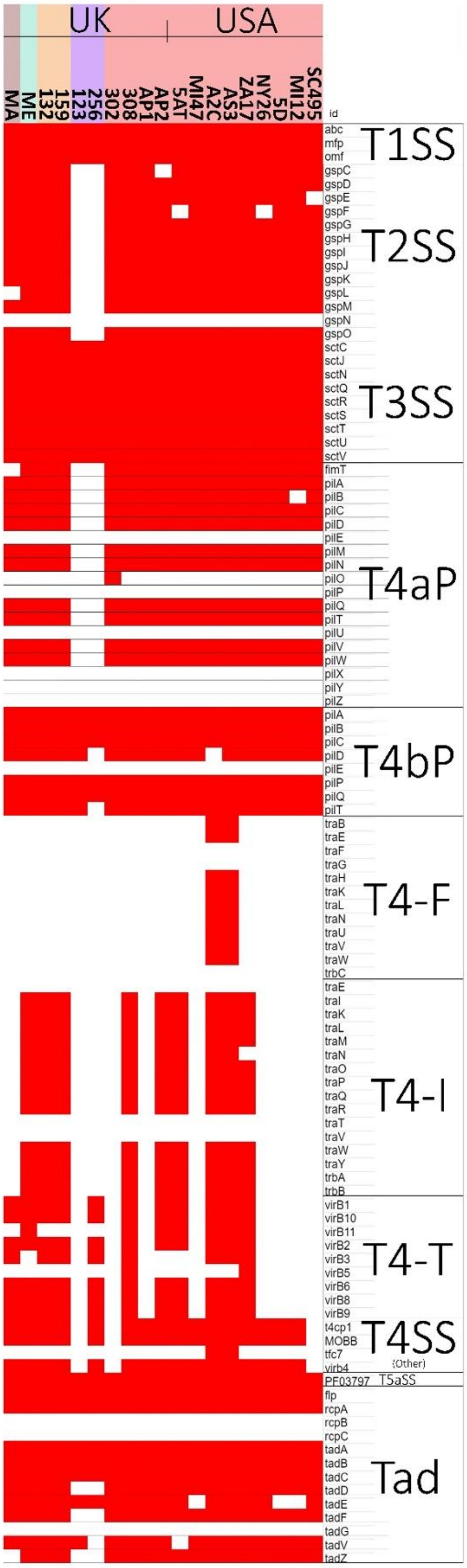


**Figure S1**. Presence (red) and absence (white) of secretion system genes in *H. defensa* genomes from three *A. pisum* biotypes - *Lotus* (orange), *Medicago* (red), and *Ononis* (purple) - plus *M. euphorbiae* (Me) and *M. artemisiae* (Ma) from the UK, and *A. pisum* strains from the USA.

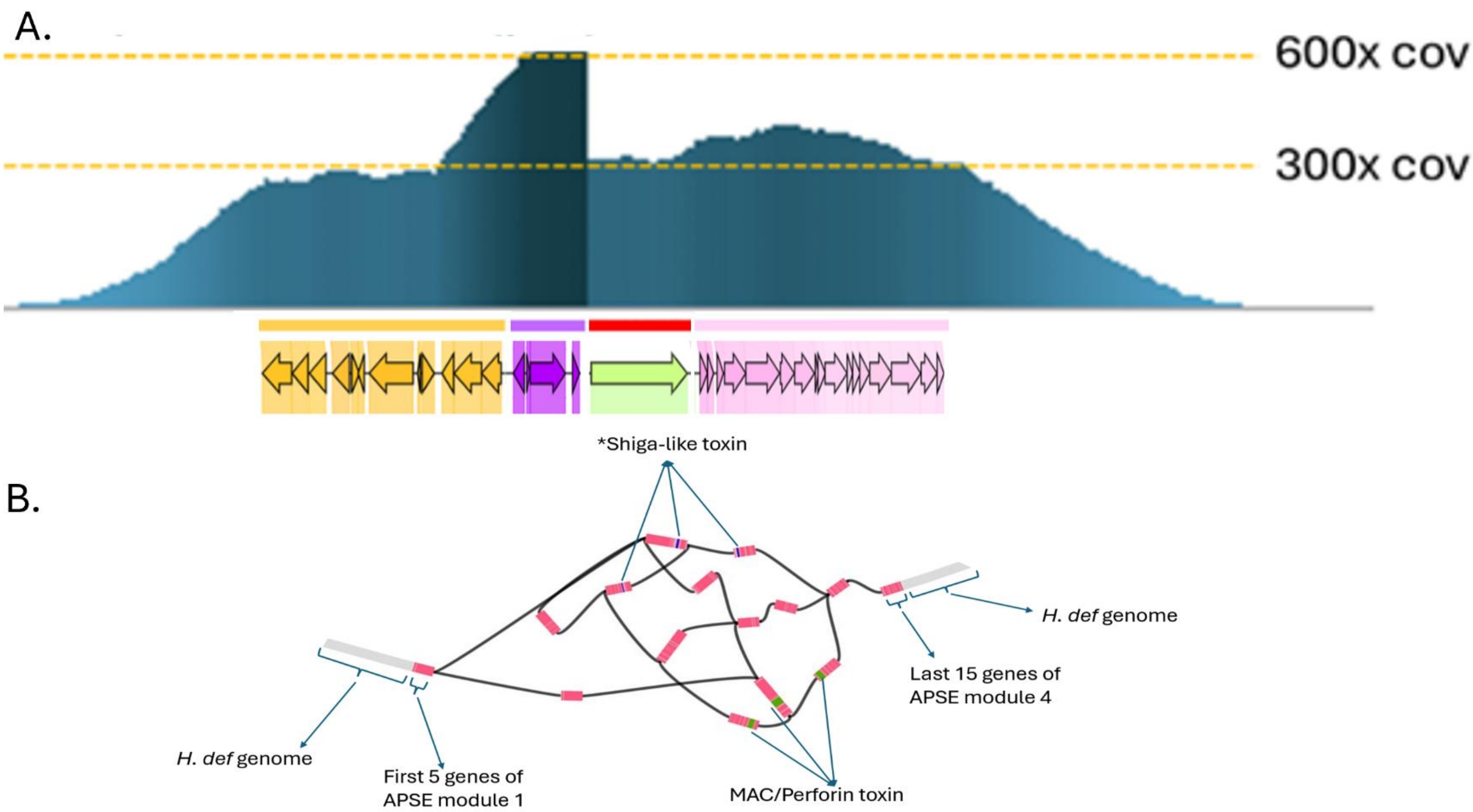


**Figure S2.** Technical evidence suggesting the presence of multiple APSE strains in the *Lotus Hamiltonella* samples (132 and 159). **A)** Coverage graph showing the number of PacBio sequencing reads successfully mapping to one reference APSE sequence. The graph shows a 300x coverage of the initial few APSE genes of module 1, at a similar coverage to the *Hamiltonella defensa* genome which was also sequenced at 300x coverage. The coverage then increases to 600x, suggesting two copies of the last few genes of module 1. The coverage of module 2 is at 600x throughout, suggesting a fully duplicated module 2. A sudden drop from 600x to 300x is seen at the border of toxin-bearing module 3, indicating that each half the APSE reads correspond to each of the two identified toxins. The results are identical when the reference APSE is exchanged for the one carrying the other toxin gene, giving each module 3 a 300x coverage from the reads generated from sequencing the same initial *Hamiltonella* sample. Coverage increases to above 300x after the toxin module, suggesting that in module 4, each APSE strain has its own few unique genes initially, whereas the last genes of module 4 seem to be shared due to the coverage drop of back to 300x. **B)** Assembly graph of the haplotype-phased unitigs in the step prior to the complete assembly of the genome sequence. From left to right, the graph shows the contiguous *Hamiltonella defensa* genome which cannot be phased into different haplotypes until reaching the APSE locus. The first 5 genes of module 1 are shared, followed by 2 haplotypes of the APSE backbone, each carrying one of the two identified toxin genes, after which the graph converges back into one haplotype carrying most of the APSE backbone genes in module 4.

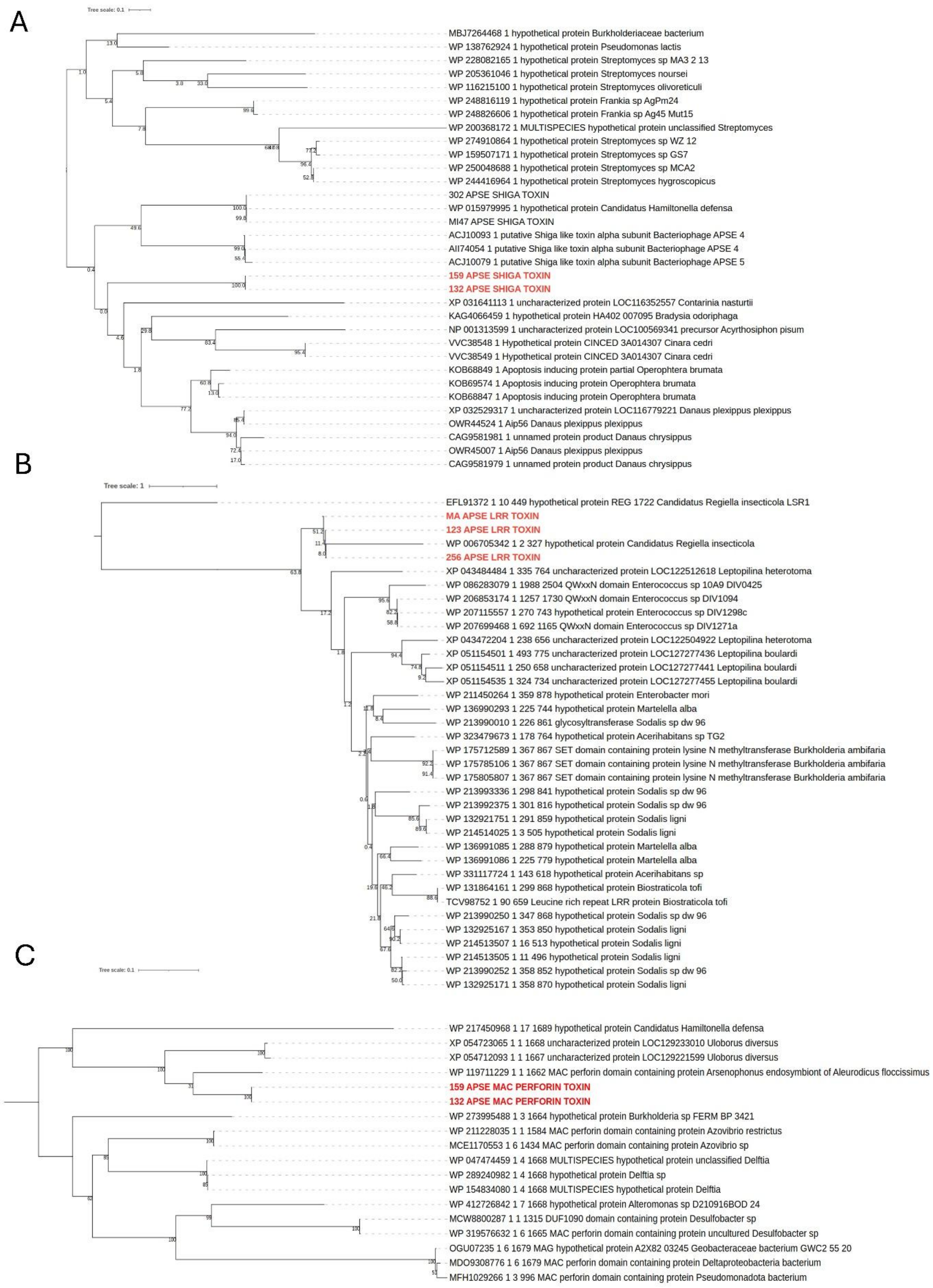
A
Tree scale: 0.1
MBJ7264468 1 hypothetical protein Burkholderiaceae bacterium
WP 138762924 1 hypothetical protein Pseudomonas lactis
WP 228082165 1 hypothetical protein Streptomyces sp MA3 2 13
WP 205361046 1 hypothetical protein Streptomyces noursei
WP 116215100 1 hypothetical protein Streptomyces olivoreticuli
WP 248816119 1 hypothetical protein Frankia sp AgPm24
WP 248826606 1 hypothetical protein Frankia sp Ag45 Mut15
WP 200368172 1 MULTISPECIES hypothetical protein unclassified Streptomyces
WP 274910864 1 hypothetical protein Streptomyces sp WZ 12
WP 159507171 1 hypothetical protein Streptomyces sp GS7
WP 250048688 1 hypothetical protein Streptomyces sp MCA2
WP 244416964 1 hypothetical protein Streptomyces hygroscopicus
302 APSE SHIGA TOXIN
WP 015979995 1 hypothetical protein Candidatus Hamiltonella defensa
MI47 APSE SHIGA TOXIN
ACJ10093 1 putative Shiga like toxin alpha subunit Bacteriophage APSE 4
AII74054 1 putative Shiga like toxin alpha subunit Bacteriophage APSE 4
ACJ10079 1 putative Shiga like toxin alpha subunit Bacteriophage APSE 5
159 APSE SHIGA TOXIN
132 APSE SHIGA TOXIN
XP 031641113 1 uncharacterized protein LOC116352557 Contarinia nasturtii
KAG4066459 1 hypothetical protein HA402 007095 Bradysia odoriphaga
NP 001313599 1 uncharacterized protein LOC100569341 precursor Acyrthosiphon pisum
VVC38548 1 Hypothetical protein CINCED 3A014307 Cinara cedri
VVC38549 1 Hypothetical protein CINCED 3A014307 Cinara cedri
KOB68849 1 Apoptosis inducing protein partial Operophtera brumata
KOB69574 1 Apoptosis inducing protein Operophtera brumata
KOB68847 1 Apoptosis inducing protein Operophtera brumata
XP 032529317 1 uncharacterized protein LOC116779221 Danaus plexippus plexippus
OWR44524 1 Aip56 Danaus plexippus plexippus
CAG9581981 1 unnamed protein product Danaus chrysippus
OWR45007 1 Aip56 Danaus plexippus plexippus
CAG9581979 1 unnamed protein product Danaus chrysippus
B
Tree scale: 1
EFL91372 1 10 449 hypothetical protein REG 1722 Candidatus Regiella insecticola LSR1
MA APSE LRR TOXIN
123 APSE LRR TOXIN
WP 006705342 1 2 327 hypothetical protein Candidatus Regiella insecticola
256 APSE LRR TOXIN
XP 043484484 1 335 764 uncharacterized protein LOC122512618 Leptopilina heterotoma
WP 086283079 1 1988 2504 QWxxN domain Enterococcus sp 10A9 DIV0425
WP 206853174 1 1257 1730 QWxxN domain Enterococcus sp DIV1094
WP 207115557 1 270 743 hypothetical protein Enterococcus sp DIV1298c
WP 207699468 1 692 1165 QWxxN domain Enterococcus sp DIV1271a
XP 043472204 1 238 656 uncharacterized protein LOC122504922 Leptopilina heterotoma
XP 051154501 1 493 775 uncharacterized protein LOC127277436 Leptopilina boulardi
XP 051154511 1 250 658 uncharacterized protein LOC127277441 Leptopilina boulardi
XP 051154535 1 324 734 uncharacterized protein LOC127277455 Leptopilina boulardi
WP 211450264 1 359 878 hypothetical protein Enterobacter mori
WP 136990293 1 225 744 hypothetical protein Martelella alba
WP 213990010 1 226 861 glycosyltransferase Sodalis sp dw 96
WP 323479673 1 178 764 hypothetical protein Acerihabitans sp TG2
WP 175712589 1 367 867 SET domain containing protein lysine N methyltransferase Burkholderia ambifaria
WP 175785106 1 367 867 SET domain containing protein lysine N methyltransferase Burkholderia ambifaria
WP 175805807 1 367 867 SET domain containing protein lysine N methyltransferase Burkholderia ambifaria
WP 213993336 1 298 841 hypothetical protein Sodalis sp dw 96
WP 213992375 1 301 816 hypothetical protein Sodalis sp dw 96
WP 132921751 1 291 859 hypothetical protein Sodalis ligni
WP 214514025 1 3 505 hypothetical protein Sodalis ligni
WP 136991085 1 288 879 hypothetical protein Martelella alba
WP 136991086 1 225 779 hypothetical protein Martelella alba
WP 331117724 1 143 618 hypothetical protein Acerihabitans sp
WP 131864161 1 299 868 hypothetical protein Biostraticola tofi
TCV98752 1 90 659 Leucine rich repeat LRR protein Biostraticola tofi
WP 213990250 1 347 868 hypothetical protein Sodalis sp dw 96
WP 132925167 1 353 850 hypothetical protein Sodalis ligni
WP 214513507 1 16 513 hypothetical protein Sodalis ligni
WP 214513505 1 11 496 hypothetical protein Sodalis ligni
WP 213990252 1 358 852 hypothetical protein Sodalis sp dw 96
WP 132925171 1 358 870 hypothetical protein Sodalis ligni
C
Tree scale: 0.1
WP 217450968 1 17 1689 hypothetical protein Candidatus Hamiltonella defensa
XP 054723065 1 1 1668 uncharacterized protein LOC129233010 Uloborus diversus
XP 054712093 1 1 1667 uncharacterized protein LOC129221599 Uloborus diversus
WP 119711229 1 1 1662 MAC perforin domain containing protein Arsenophonus endosymbiont of Aleurodicus floccissimus
159 APSE MAC PERFORIN TOXIN
132 APSE MAC PERFORIN TOXIN
WP 273995488 1 3 1664 hypothetical protein Burkholderia sp FERM BP 3421
WP 211228035 1 1 1584 MAC perforin domain containing protein Azovibrio restrictus
MCE1170553 1 6 1434 MAC perforin domain containing protein Azovibrio sp
WP 047474459 1 4 1668 MULTISPECIES hypothetical protein unclassified Delftia
WP 289240982 1 4 1668 hypothetical protein Delftia sp
WP 154834080 1 4 1668 MULTISPECIES hypothetical protein Delftia
WP 412726842 1 7 1668 hypothetical protein Alteromonas sp D210916BOD 24
MCW8800287 1 1 1315 DUF1090 domain containing protein Desulfobacter sp
WP 319576632 1 6 1665 MAC perforin domain containing protein uncultured Desulfobacter sp
OGU07235 1 6 1679 MAG hypothetical protein A2X82 03245 Geobacteraceae bacterium GWC2 55 20
MDO9308776 1 6 1679 MAC perforin domain containing protein Deltaproteobacteria bacterium
MFH1029266 1 3 996 MAC perforin domain containing protein Pseudomonadota bacterium

**Figure S3**. Maximum likelihood amino acid phylogenies of the newly identified APSE toxin genes highlighted in red text. Trees were rooted at the midpoint to allow for the phylogenetic placement of each toxin gene among closest related protein coding sequences on Genbank. **A)** Phylogeny of a new shiga-like toxin gene found in the APSE bacteriophages of *Hamiltonella* from *Lotus* biotype (strains 132, 159) pea aphids. The gene is shown to be adjacent to a clade of previously identified Shiga-like toxins from APSE phages. **B)** Phylogeny of a new Leucine-rich-repeat domain containing putative toxin gene found in the APSE bacteriophages of *Hamiltonella* from *Ononis* biotype (strains 123, 256) pea aphids and *Hamiltonella* (strain MA) from the *M. artemisiae* aphid. Low bootstrap values imply the lack of closely related genes. **C)** Phylogeny of a new MAC/Perforin domain containing putative toxin found in the APSE bacteriophages of *Hamiltonella* from *Lotus* biotype (strains 132, 159).

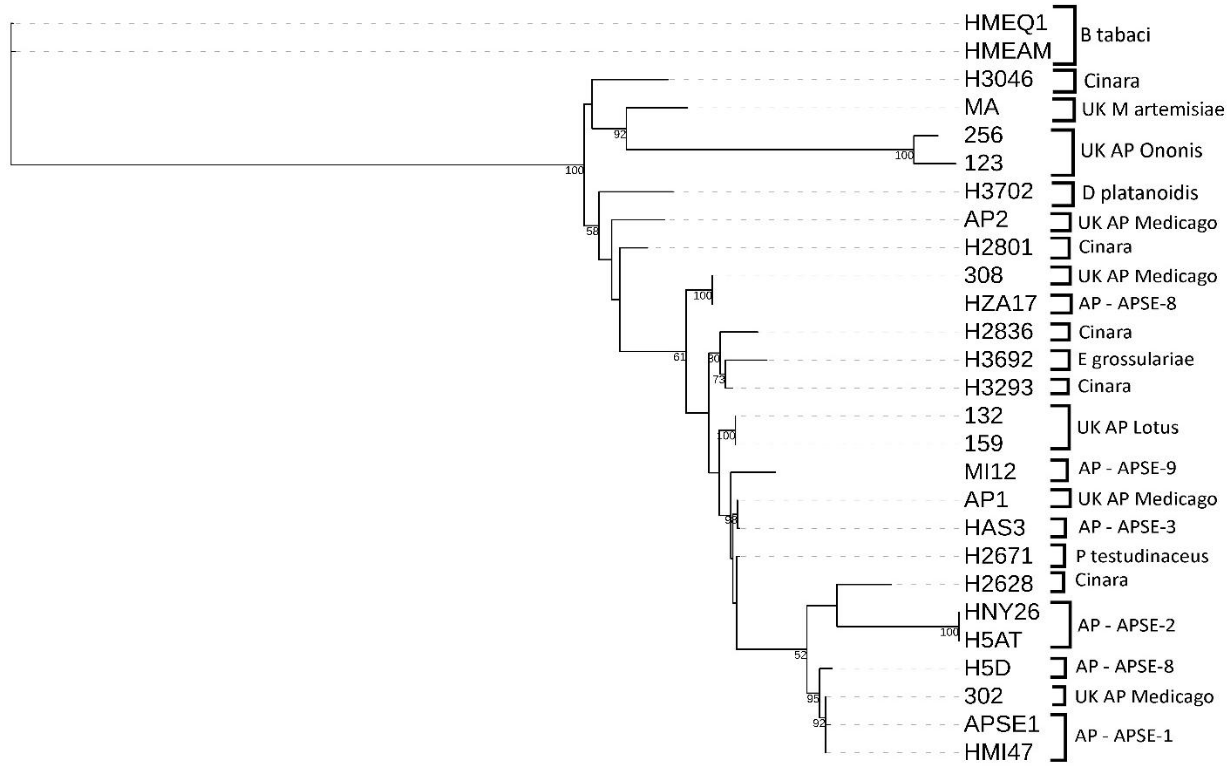


**Figure S4**. Maximum-Likelihood core gene phylogeny of the orthologous genes found in all APSE backbones built with 500 bootstraps. Tips of phylogeny are annotated with original sample names of their respective *Hamiltonella defensa* host genomic sources, followed by the parent insect names in brackets.

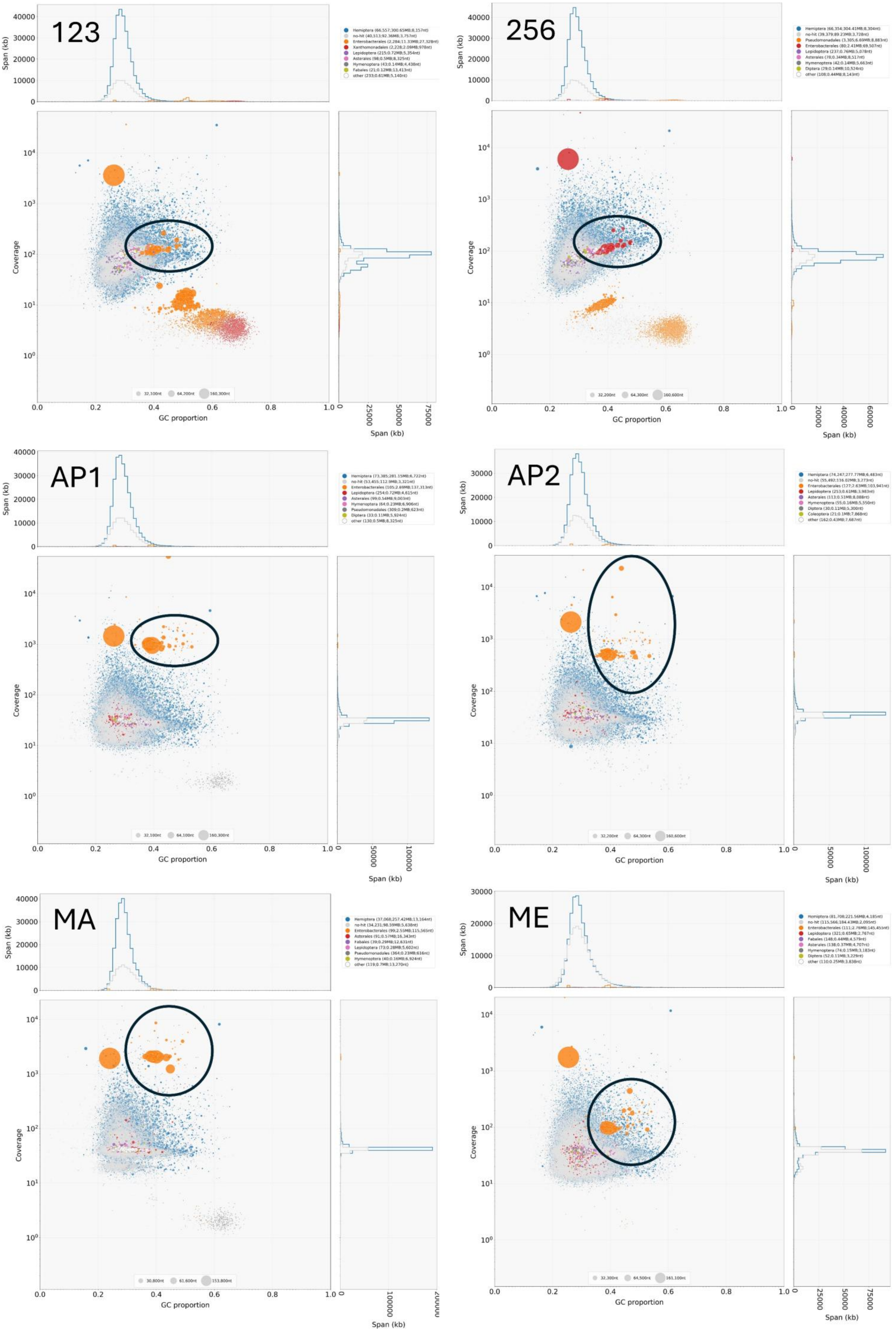
123
256
AP1
AP2
MA
ME
Span (kb)
Coverage
GC proportion

**Figure S5**. Blobplots of the Illumina sequenced aphid-*Hamiltonella* samples. Initial assemblies of entire metagenomes showing the distribution of sequence coverage on the Y axis, GC% on the X axis, taxonomic ID through different colours of blobs, and sizes of initial contigs by the size of the blobs. Blobs corresponding to *Hamiltonella* genomes are circled.
